# Purifying selection and drift, not life history or RNAi, determine transposable element evolution

**DOI:** 10.1101/034884

**Authors:** Amir Szitenberg, Soyeon Cha, Charles H. Opperman, David M. Bird, Mark Blaxter, David H. Lunt

## Abstract

Transposable elements (TEs) are a major source of genome variation across the branches of life. Although TEs may occasionally play an adaptive role in their host’s genome, they are much more often deleterious, and purifying selection is thus an important factor controlling genomic TE loads. In contrast, life history and genomic characteristics such as mating system, parasitism, GC content, and RNAi pathways, have been suggested to account for the startling disparity of TE loads in different species. Previous studies of fungal, plant, and animal genomes have reported conflicting results regarding the direction in which these genomic features drive TE evolution. Many of these studies have had limited power because they studied taxonomically narrow systems, comparing only a limited number of phylogenetically independent contrasts, and did not address long term effects on TE evolution. Here we explicitly test the long term determinants of TE evolution by comparing 42 nematode genomes that span over 500 million years of diversification, and include numerous transitions between life history states and RNAi pathways. We have analysed the reconstructed TE loads of ancestors through the Nematoda phylogeny to account for correlation with GC content and transitions in TE evolutionary models. We also analysed the effect of transitions in life history characteristics and RNAi using ANOVA of phylogenetically independent contrasts. We show that purifying selection against TEs is the dominant force throughout the evolutionary history of Nematoda, as indicated by reconstructed ancestral TE loads, and that strong stochastic Ornstein-Uhlenbeck processes are the underlying models which best explain TE diversification among extant species. In contrast we found no evidence that life history or RNAi variations have a significant influence upon genomic TE load across extended periods of evolutionary history. We suggest that these are largely inconsequential to the large differences in TE content observed between genomes and only by these large-scale comparisons can we distinguish long term and persistent effects from transient effects or misleading random changes.

## Introduction

Transposable elements (TEs) are mobile genetic entities found in the genomes of organisms across diverse branches of life, and which are a major source of genetic variation [1–3]. TEs comprise approximately two thirds of the human genome [4], and in some plants and animals they may account for 85% of the genome [5,6]. In stark contrast, other plant and animal genomes contain only 1–3% TE-derived sequence within their typically much smaller genomes [7,8]. The mechanisms that create this variability are not fully understood.

TE insertions are a significant source of deleterious mutation causing gene disruption [1,9], double-strand breaks [10, 11], ectopic recombination [12], gene expression change [13], and other types of mutagenesis [14]. In humans, deleterious TE activity contributes to approximately 0.3% of genetic disease [15, 16]. Some TE insertions, however, have only weak deleterious effects, increasing their likelihood of survival and expansion [17–22], Given sufficient time, a small proportion of these may be co-opted for protein-coding or regulatory functions by the host genome, and thus become very important components of organismal evolution [13,23,24]. Despite being a key player in organismal evolution, the evolutionary forces determining the TE composition in genomes are far from clear. We have selected the phylum Nematoda, for its phylogenetic diversity of available genomes, as a system in which to investigate TE variation in a phylogenetically-controlled design. While other studies have examined the correspondence between life history or other traits with TE evolution [25–29], these often muster relatively few phylogenetically independent contrasts, and a relatively recent evolutionary time scale. Examining evolutionary events across the entire phylum Nematoda gives a broad perspective where the balance of evolutionary forces will have had time to work.

Substantial efforts to characterise the forces and processes shaping genome evolution have given rise to explanations for the divergence in TE loads among species, including the effects of mating system and recombination, life history, genome GC content, and transposon removal systems such as RNAi. These factors influence TEs both directly, by affecting their possibility for spread or removal, and indirectly, by modifying the effective population size and probability of fixation [cf. 30, inf.]. The effects of mating system and recombination have been much discussed, with conflicting predictions for either an increase [31, 32] or decrease [33–37] in TE loads in selfing species. Duret, et al. [38] found that non-recombining genomic regions are less TE rich than recombining regions in *Caenorhabditis elegans*, when considering DNA TEs. Also in *Caenorhabditis*, Cutter, et al. [26] predicted lower TE loads in selfing compared to outcrossing species. In contrast, in *Drosophila* TE spread was positively associated with recombination [28,29]. A mating system effect on genome size (and thus likely TE load), was reported in plants [39–41], but subsequent studies accounting for phylogenetic associations in the data did not recover these effects [27,42,43]. Analysis of the evolution of TE loads in the Nematoda, where several independent shifts in mating system have occurred (Fig. 1), may aid in resolution of these issues.

**Fig 1.**
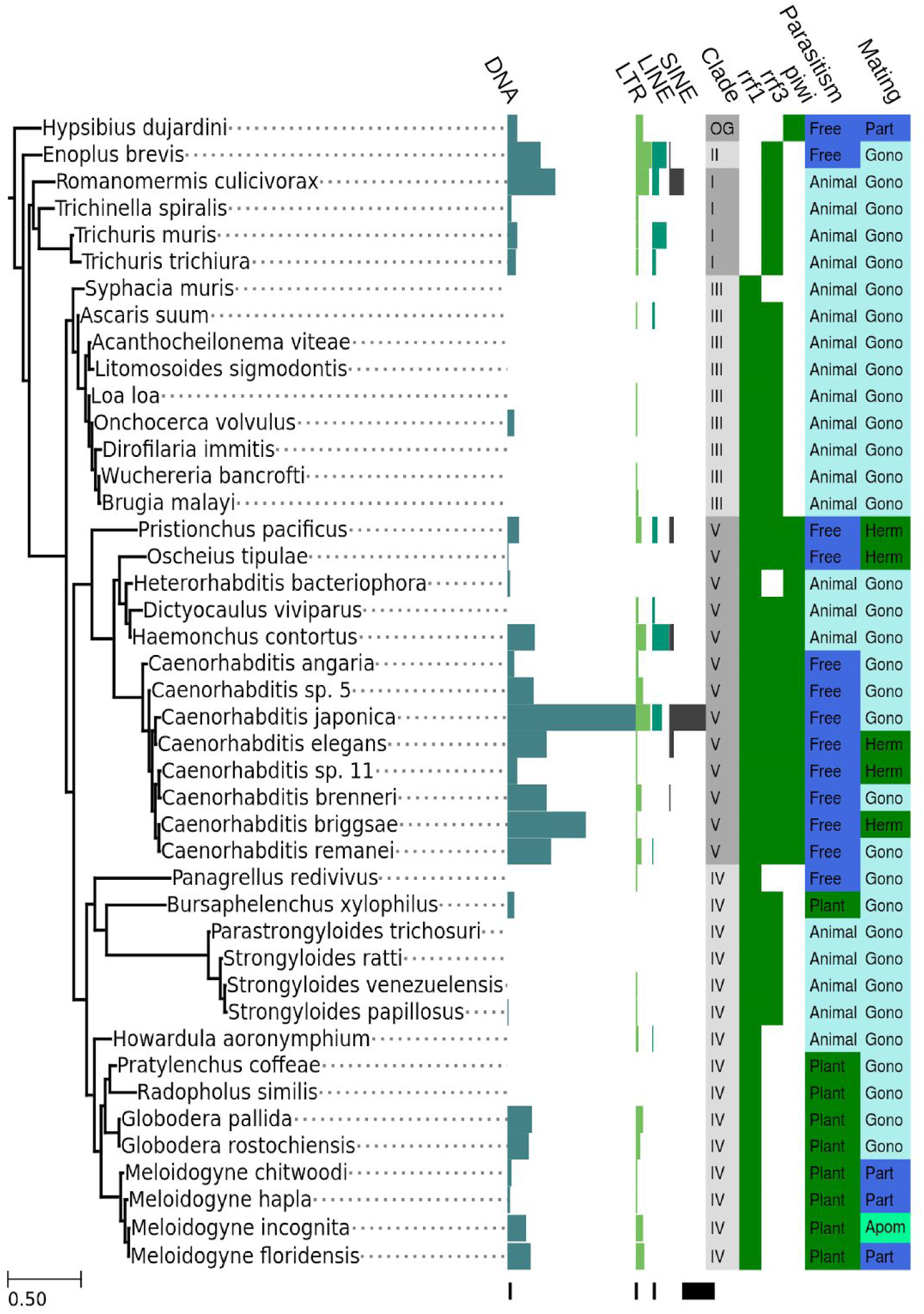
TE loads in Nematoda by class. SSU-rRNA phylogenetic tree of Nematoda with TE load information by class. The columns represent (left to right) DNA, LTR, LINE and SINE element loads (numerical values are given in S2 Table), the phylogenetic clade *sensu [78]*, presence or absence of RNAi pathway proteins (RRF1, RRF3 and PIWI), parasitism (animal parasite, plant parasite, or free living), and mating system (parthenogenic, gonochoristic, hermaphroditic or apomictic). Black scales at the bottom of each bar-chart represent 2500 TEs. Sources for life history information are in S1 Table.

Adoption of a parasitic lifestyle can reduce the effective population size, and thus the effectiveness of recombination. Parasites may be subdivided into infrapopulations within hosts, and this population subdivision reduces the effective population size compared to free-living species [44]. Increased TE counts were found in ectoparasitic *Amanita* fungi compared to free living *Amanita* species [25], where the authors suggested the effective population size effects of parasitism as a cause for the difference. As Nematoda contain several independent transitions to parasitism, this hypothesis can also be further tested (Fig. 1).

Genome nucleotide bias (GC content) has been shown to influence a wide variety of cellular processes, and especially the rates and patterns of molecular evolution. These effects include tRNA abundance and codon usage [45–48], mutational patterns [49,50], gene expression [51–56], protein and RNA structure and composition [57–63], and translational efficiency [64]. The tight integration of TEs with cellular processes will mean that they will also be affected by differential nucleotide biases, as has been examined by Hellen and Brookfield [22], who demonstrated the accumulation and persistence of human *Alu* elements to be favoured in GC-rich regions. Again, diversity in GC content across nematode genomes offers power to detect the effects of GC on TE load evolution.

The host genome is engaged in defending itself against TE insertions, with RNA interference (RNAi) pathways a key cellular processes silencing TEs in eukaryotes [65–72]. In nematodes a variety of mechanisms of TE silencing have been characterised at the molecular level [73–76], with different pathways operating in different clades (Fig 1). This variation permits examination of the role of alternate TE silencing pathways in explaining genome-wide TE loads.

If differences in TEs between lineages are not determined by processes such as mating system or life history then we are drawn back to our null model of genome evolution, one which is shaped by non-deterministic processes such as mutation and drift [77]. The importance of non-deterministic processes in shaping TE evolution has been long recognized with Charlesworth and Charlesworth [30] hypothesising that the efficiency of selection and TE silencing depends on the effective population size. Here we conduct correlation and ANOVA tests of the deterministic forces previously proposed to affect TE evolution, with phylogenetically independent contrasts of TE counts in species from across the phylogenetic diversity of Nematoda [78] as the dependant variable. We find no evidence that any of the deterministic forces significantly explain differences in TEs among lineages. While we demonstrate that TE loads are shaped by patterns of purifying selection in the long term, our data strongly suggest that stochastic changes are the major genome-wide determinant of TE diversity.

## Results

### TE loads in Nematoda

To test the effect of the mating system, parasitic lifestyle, GC content, RNAi and transposition mechanism on TE evolution, TEs were identified and classified in 43 genome assemblies representing the five major nematode lineages and the tardigrade *Hypsibius dujardini* (Fig 1, S1 Table, S1 Methods, sections 1 to 7). TE loads did not correlate with genome assembly quality (represented by N50 values), assembly size or predicted genome size (S1 Methods, section 1). High TE loads have a patchy distribution among species in Nematoda, with hotspots observed in the Dorylaimia (Clade I of Blaxter et al. [78]) and Enoplia (Clade II), in Rhabditina (Clade V), and in the Tylenchomorpha genera *Meloidogyne* and *Globodera* (part of Clade IV) (Fig 1). DNA elements were usually the most abundant, followed by LTR elements, while LINE and SINE elements were quite scarce (Fig 1). When classes were broken down into families (Fig 2, S2 Table, S1 Figure), a large proportion of the variation among species, for ‘cut and paste’ DNA elements, was contributed by variation in loads of *TcMar* element families, which are scarce in Dorylaimida (Clade I) and abundant in Rhabditina (Clade V). *hAT* families followed a similar pattern, but with less extreme differences among species. *Onchocerca volvulus* (Spirurina; Clade III) had high loads of Helitron elements (5372 copies), and hardly any other TEs, a very different pattern from its relatives in Clade III. Among LTR superfamilies, *Gypsy* elements predominated, with *Copia* and *Pao* elements also prevalent, though a large proportion of the elements were unclassified. The predominant LINE elements were *Penelope* and RTE. SINE elements, although more abundant in a few Rhabditina (Clade V) species than in others, were generally scarce (< 500 in most species, S2 Table).

**Fig 2.**
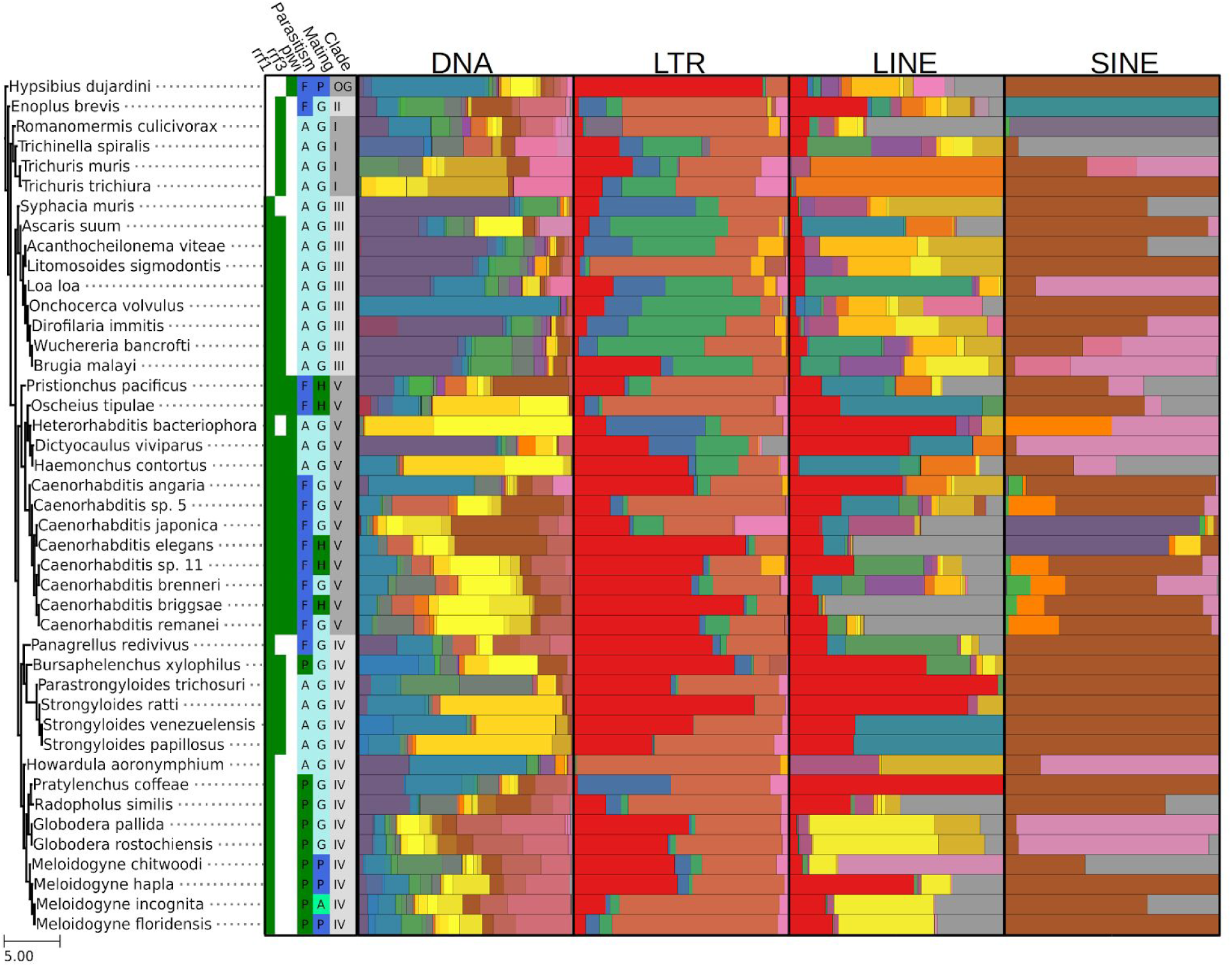
TE loads in Nematoda by superfamily. SSU-rRNA phylogenetic tree of Nematoda with TE loads information by superfamily. The columns represent (left to right) the presence or absence of RRF1, RRF3 and PIWI RNAi pathway proteins, parasitism (A-animal parasite, P-plant parasite, or F-free living), and mating system (P-parthenogen, G-gonochoric, H-hermaphroditic or A-apomictic), the phylogenetic clade *sensu [78]* and the proportions of DNA, LTR, LINE and SINE element superfamilies within each of the classes (numerical values in S2 Table, colour key in S1 Figure).

### Phylogenetic signal in TE load

According to our null hypothesis, TE load evolves neutrally and change (in rates and patterns) is expected to be congruent with the topology and branch lengths of the species tree. This can be assessed via phylogenetic transformations of observed TE loads [79]. To account for phylogenetic uncertainty while computing such transformations, we generated a Bayesian posterior distribution of SSU-rRNA phylogenetic species trees. Tree transformation values of the TE counts were computed with each of the trees in the posterior distribution, and for each of the TE classes (DNA, LTR, LINE and SINE; Fig 3A). Transformation value distributions were also computed for each superfamily (S2 Figure), within each class of TEs, and the median value of each superfamily was recorded across the superfamilies in a given class (Fig 3B). We did not include SINE element superfamily medians, since SINE elements were too sparse to compute a meaningful distribution (Fig 3B).

**Fig 3.**
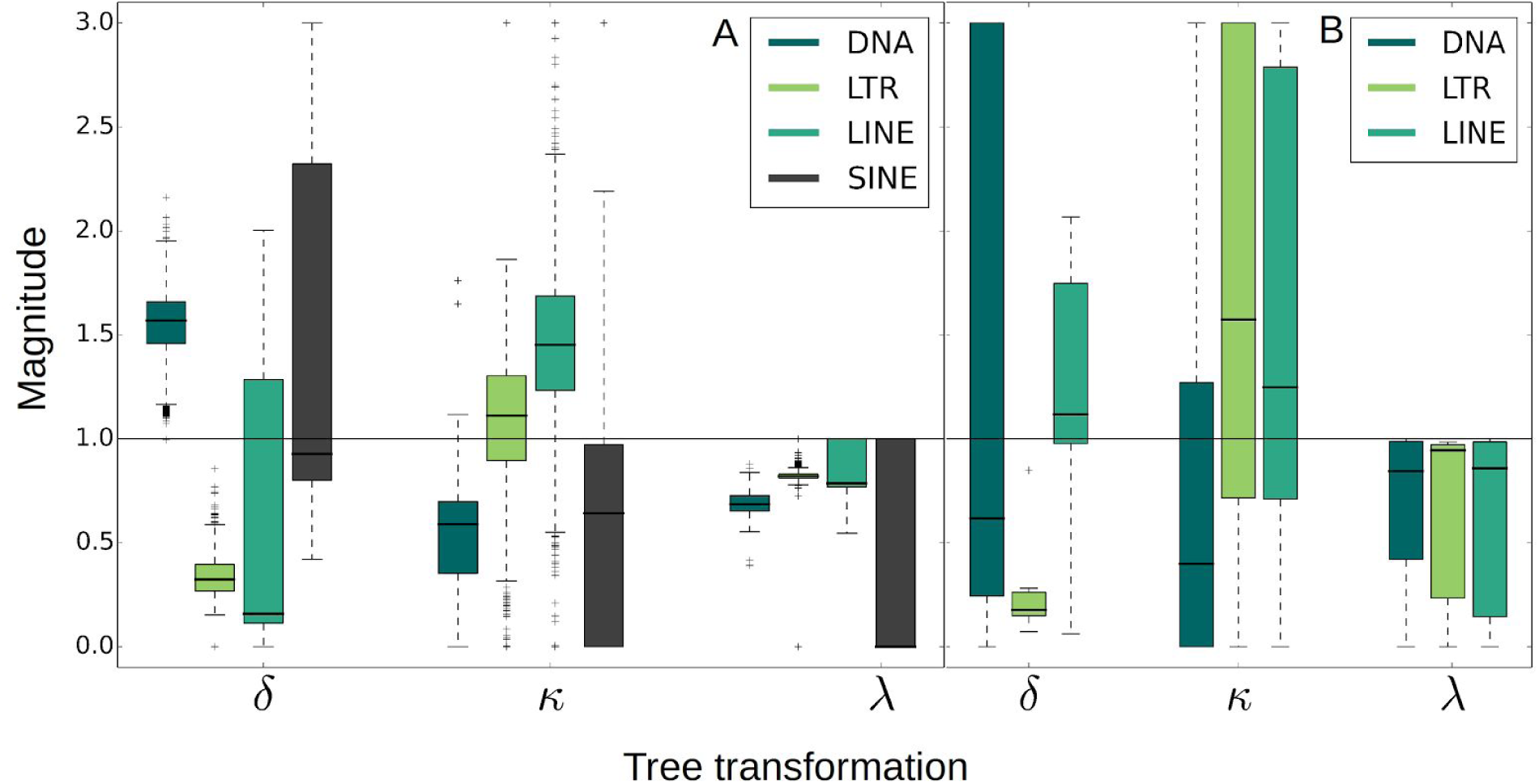
Phylogenetic transformations of TE loads. The *δ, κ*, and *λ* transformations of TE loads, representing the fit between the TE loads and the tree’s topology (*λ*), branch-lengths (*κ*) and root-tip distance (*δ*). (A) The distribution of transformation values across the posterior distribution of most likely phylogenetic trees for each element class (DNA, LTR, LINE and SINE). (B) The distribution of median transformation values of each superfamily of elements within each of the classes. Only superfamilies where the distance between the first and third quartiles was smaller than 0.2 for *λ* and smaller than 0.5 for *κ* and *δ* are included (i.e., superfamilies with an unresolved transformation value are excluded). SINE elements are not shown because the distributions cover the whole range of values. Per-superfamily distribution of the λ, *κ* and *δ* transformations across the posterior distribution of trees is shown in S2 Figure.

The *λ* transformation [79] p rovides an estimate of the correlation between the TE quantities and the topology of a tree with *λ* = 1 indicating a strong correlation. At the class level, DNA, LTR and LINE element load variations are strongly correlated with the species phylogenetic relationships (λ > 0.5; Fig 3A). For many superfamilies the median *λ* was greater than 0.5, indicating that the correlation with the phylogeny is a general characteristic of TEs, and not only a feature of a few large superfamilies (Fig 3B). For SINE elements, in part due to their low abundance and phylogenetic uncertainty, this correlation was not recovered. A second phylogenetic transformation *κ*, provides an estimate of the correspondence between the branch lengths and the rate of change of a trait [79]. *κ* > 1 indicates a higher rate of change in longer branches, *κ* = 1 indicates that the rate of change of the trait conforms with the general evolutionary rate, and *κ* < 1 indicates that the trait is more conserved than expected from neutrality. The *κ* value distribution for nematode DNA TE loads showed that DNA TE evolution depends less on the organismal evolutionary rate than other TE classes, at the class level (*κ* < 1; Fig 3A). The pattern persisted for most superfamilies when considering *κ* median values at the DNA element superfamily level (Fig 3B). Lastly, the *δ* transformation estimates the tree depth at which non-neutral evolutionary events occurred, where *δ* < 1 suggests ancient events and *δ* > 1 indicates that the trait diversified recently. For DNA elements, *δ* was greater than 1, indicating that recent events explain their current TE load patterns, while for LTR elements, *δ* was less than 1, indicating that ancient events explain their load patterns. *δ* was not determined for LINE and SINE elements due to phylogenetic uncertainty. Only for LTR elements did these findings persist when the median *δ* values of individual superfamilies were considered (Fig 3B), where all of the LTR superfamilies underwent important early events (median *δ <* 0.3).

### The effect of life cycle, RNAi pathway, and genome GC content variation on TE evolution

Primary literature was surveyed in order to determine the mating system of each species and to identify parasites of plants and animals (S1 Table). Key proteins involved in RNA silencing of transposons (RRF1, RRF3 and PIWI) were identified in the genome assembly data using reference sequences (from [76], S1 Methods, section 8), and genome assembly N50, span and GC content were calculated. The reproductive mode, parasitic status and RNAi pathway for each nematode species is summarized in Fig 1 and S1 Table. The presence and absence of RNAi pathway proteins for the most part conformed with the predictions made by Sarkies et al. [76], with a few exceptions. *Syphacia muris*, (Oxyuridomorpha, Spirurina in Clade III), lacks the expected RdRP RRF3 protein that is found in other Spirurina species. Since the genome assembly has high N50 values (60,730 bp), and much supporting transcriptome data (S1 Methods, section 8.9), it is highly likely that this species lacks RRF3 (or possess a very divergent RRF3 orthologue). The *Heterorhabditis bacteriophora* (Rhabditomorpha; Clade V) genome lacked an RRF3 locus although RRF3 is expected in Rhabditomorpha species [76]. Given the relatively high quality of the *H. bacteriophora* genome assembly (N50 of 33,765 bp), RRF3 is again likely absent (or very divergent) in this species. No RRF3 were found in any of the 9 Tylenchomorpha species (Clade IV), regardless of their N50 values (3,348 bp to- 121,687 bp), in keeping with expectations [76].

For each TE class and superfamily, we tested the effect of mating system, parasitic lifestyle, and variation of RNAi pathways on TE loads at terminal nodes using an ANOVA of phylogenetically independent contrasts. Only a few of the tests addressing mating system variations were significant. In the DNA element ISC1316 and the YR/Ngaro elements, we found a significant effect of mating system (p-values 0.013 and 0.005 respectively) when species were classified into three mating system types (gonochoristic, apomictic and facultative). RNAi pathway protein presence did not emerge as a significant factor in TE evolution, and neither did parasitism. The number of significant ANOVA test results was 0.98% of all the ANOVA tests conducted, and these were non-significant with a Holm–Bonferroni correction [80]. We also explored the correlation between genome assembly GC contents and TE loads (S1 Methods section 10.25) and found a weak and inverse correlation (r ≅ −0.3, p-value = 0.03) for the superfamily DNA/Ginger2, and also a weak correlation (r ≅ 0.4, p-value = 0.02) for the superfamily LINE/Penelope, which become non-significant with a Holm–Bonferroni correction [80].

### Changes of TE loads at ancestral nodes

To understand long term processes in TE evolution, we reconstructed the TE loads for each element superfamily at each node in the Nematoda phylogeny, and derived the median change in TE loads at each node compared to its ancestor (Fig 4). For all the four TE classes (DNA, LTR, LINE, and SINE) the evolutionary process was characterized by a trend towards contraction of TE loads, with only very few events of stable expansions, except for shallow nodes where the nature of change was less predictable. Purifying selection in deep nodes appeared to have been more constant for LTR elements than other classes (Fig 4B), in agreement with the *δ* value in this class (Fig 3), and LTR elements were also the most dynamic in shallow nodes, with shallow expansion hotspots within Onchocercidae (Clade III), Strongylida and *Caenorhabditis* (Clade V), and *Stmngyloides, Globodera* and *Meloidogyne* (CladeIV). Other classes (Fig 4A, C and D), also showed recent expansions, but only in a subset of these taxa.

**Fig 4.**
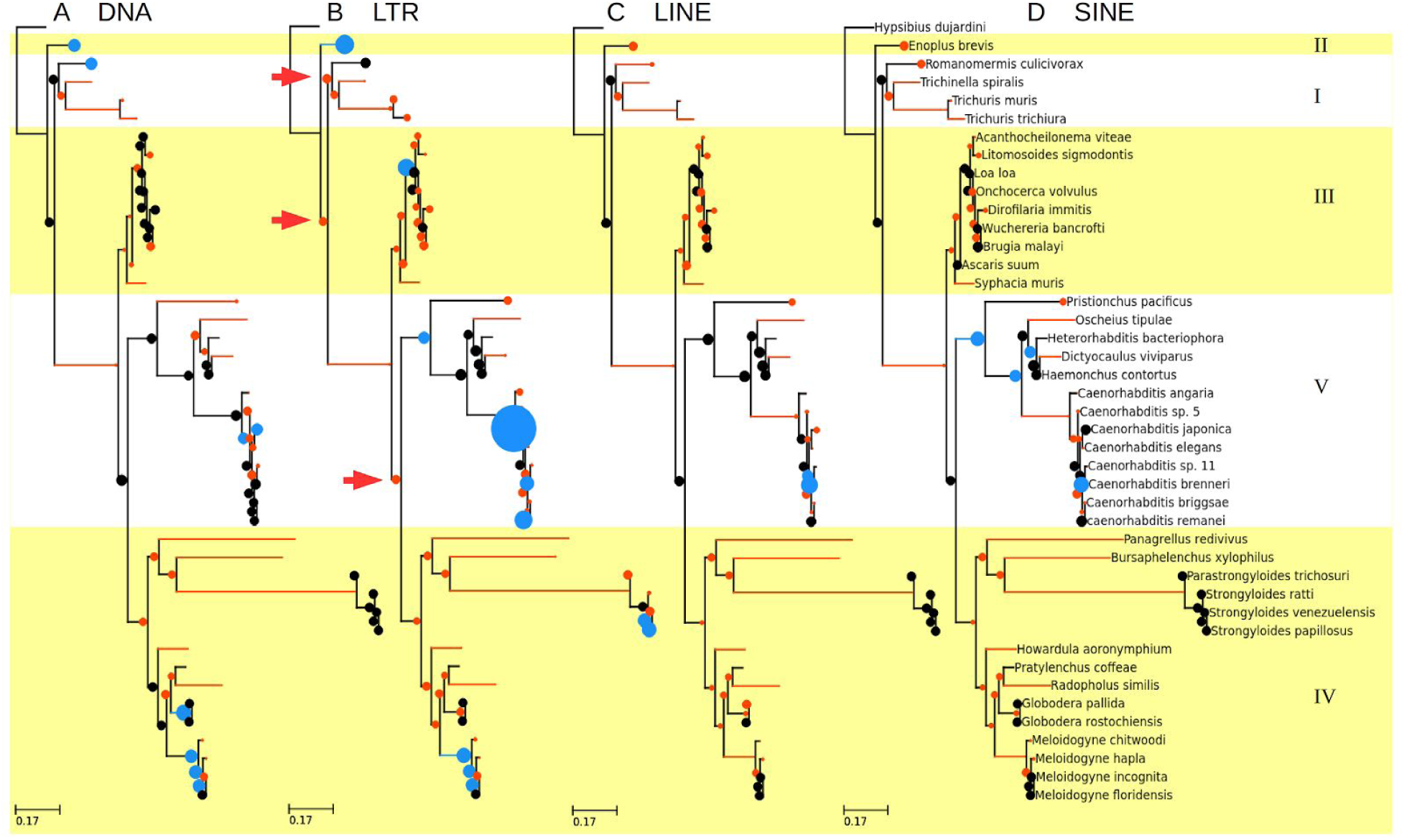
Median TE load change at ancestral and terminal nodes. The median load change of DNA (A), LTR (B), LINE (C), and SINE (D) superfamilies. Ancestral states were reconstructed for each superfamily. Then, the proportion of change, compared to the ancestral node, was computed for each superfamily, at each node. The median change proportions are presented for each class. Blue nodes represent an increase compared to the ancestral nodes, with larger nodes representing a greater increase. Orange nodes represent a decrease compared to the ancestor, with smaller nodes representing a greater decrease. Long branches (0.06 or longer) along which at least 50% change in TE loads has occurred are blue or orange to indicate a decrease or increase of TE median loads. Red arrows (B) indicate deep nodes in which a reduction in LTR elements has occurred, but not in other TE classes.

### Detection of adaptive processes and convergent evolution

To identify adaptive processes in the evolution of TEs, we tested the fit of the TE loads with the Ornstein Uhlenbeck (OU) model, using BayesTraits 2 [81]. Given the high *λ* values characterising TE evolution in Nematoda we predicted high power to detect α, the selection strength parameter in the OU process (Fig 5A). Assuming no stochastic interference, α values greater than 1 were significantly detected in fourteen, seven, three, and one superfamilies from the classes of DNA, LINE, LTR and SINE elements respectively (S3 Figure, S1 Methods, section 10.18). These families were analysed for convergent evolution, fitting the most likely extended model, also allowing shifts in the selective optimum (*θ*) as well as stochastic change (σ^2^). Convergent evolution (i.e., polyphyletic lineages possessing the same selective optimum *θ*) was detected for all these elements. However, shifts in *θ* were only identified in terminal or otherwise shallow nodes, with the exception of a *θ* increase for two LTR element superfamilies at the base of the Rhabditina (Clade V), and never coincided with shifts in mating system, parasitism, or RNAi pathways (S4 Figure). Moreover, the *σ^2^* (drift) values were overwhelmingly higher than α (selection) for most of the superfamilies, as the example shown in Fig 5B, illustrating the stochasticity of the evolutionary trajectories of TE loads.

**Fig 5.**
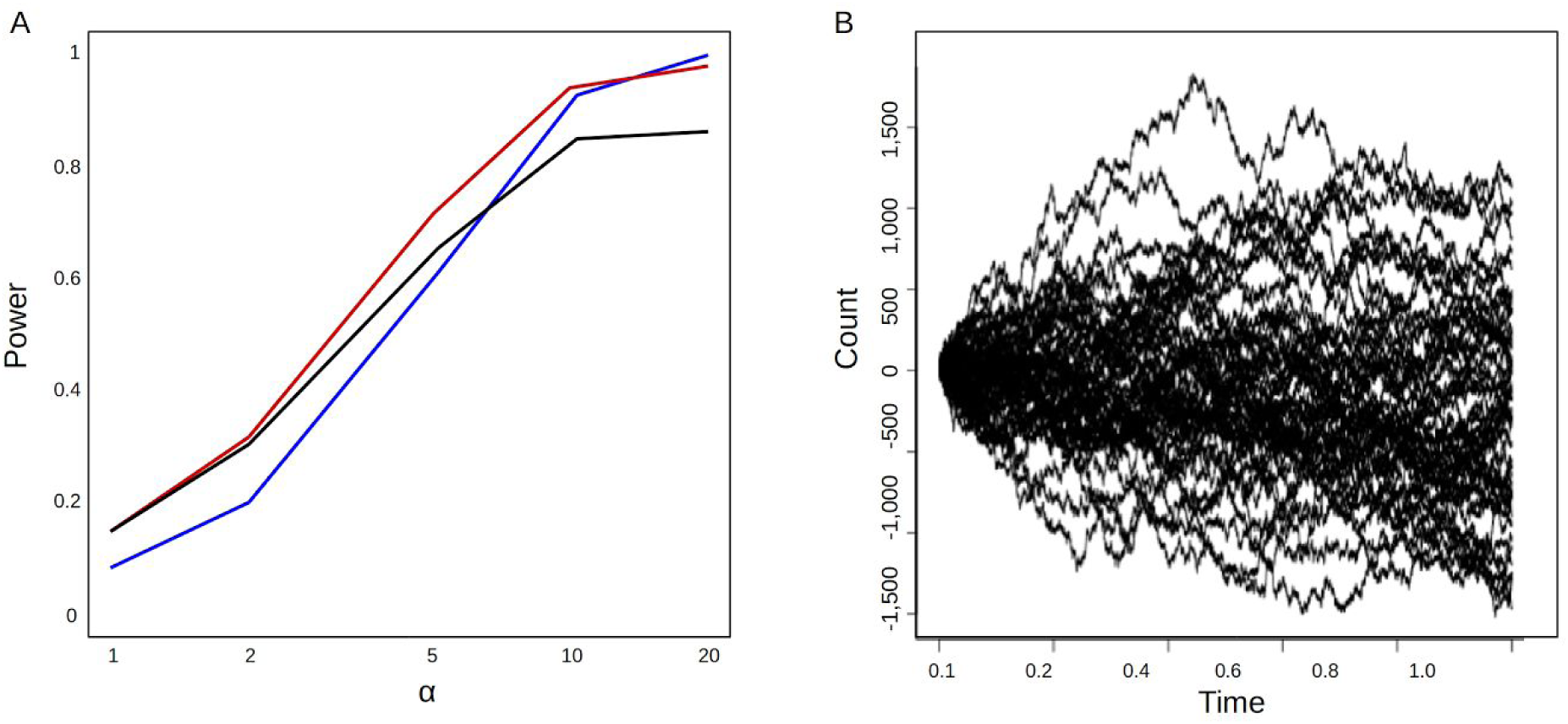
OU-model fitting to detect selection. Power to detect the selection strength parameter alpha (A), under a gamma transformation value of 0.5 (black), 0.8 (red) and 1 (blue), and simulations of the evolutionary trajectory of the DNA/TcMar-Tc2 TE superfamily loads (B) under the OU model fitted to this superfamily (*σ^2^=* 1^*^10^9^, α=4^*^10^5^ for 10^5^ generations and 50 replications).

## Discussion

The common ancestor of Nematoda dates back to the Cambrian radiation [82], 550 million years ago, and thus the genome sequences of nematodes that have become available in the last decade provide a unique opportunity for comparative genomics analyses of the long term forces shaping evolution. This contrasts with most previous studies which were only able to analyse recent periods [25–29, e.g., 38, 40, but see 42, 43, 83, 84]. We present analyses of the long term evolution of TEs, exploring the roles and importances of multiple deterministic forces in a phylogenetic design. Our results establish purifying selection as the sole effective long term deterministic force. We also find that recent diversification in TE loads is independent of GC content, life history, and RNAi, and is best understood as a stochastic process.

### Purifying selection prevails in TE evolution

We find that a model of purifying selection best explains the evolution of transposable elements, with LTR elements in particular having been purified in ancestral nodes of the Nematoda phylogeny. Furthermore, long branches in the phylogeny, including terminal ones, often coincide with reduction of at least 50% in TE loads (Fig 4) and almost never with an increase. This reveals that purifying selection prevails in the long run, over forces and events that might increase or preserve TE loads temporarily. The co-occurrence of increased purifying selection of LTR elements with their increased expansions in terminal nodes, suggests that, on average, LTR element loads have a tendency to increase faster than other elements. LTR elements are therefore more likely to have deleterious effects and are more exposed to purifying selection, as suggested previously [14,85].

The increased purifying selection experienced by LTR elements could result from either their indiscriminate targeting of genic regions [21,86] or from their role in induction of increased ectopic recombination [32]. It may be that LTR elements have not been able to evolve to efficiently target non-genic regions of the genome [87,88]. One signature of ectopic recombination as a driver of purifying selection on LTR TEs would be an inverse correlation between the median sequence length of TE families and their loads in a given species [89], but we did not detect such inverse correlation (S1 Methods, section 11).

### Long term GC content variation does not determine TE loads

Genome GC content can change gradually along the phylogenetic tree. We therefore used an analytic procedure that accounts for the ancestral character states of both TE load and GC content traits and tested for correlation between the two through the evolutionary history of nematodes. In humans, purifying selection against TE loss in gene rich regions of the genome is the main driver of variability in *Alu* element loads between GC-rich and -poor genomic regions [22,90]. However, GC variation within a genome does not explain TE load differences between species with different whole-genome GC contents as we did not find substantial correlation between the TE loads and the GC content of the nematode genome assemblies. While the local GC content may indeed influence the number of insertions fixed in a given locus, it is not a limiting factor on TE loads in the genome as a whole.

### Recent variation in RNAi pathways and life history is not a predictor of TE evolution

The strength of purifying selection on TE loads appears to be independent of recent shifts in the species life history of RNAi pathways involved in TE silencing. Less then one percent of the ANOVA tests, examining the effect of RNAi, parasitism and mating system on TE loads, suggested significant associations between traits and TE loads, well within the limits of an acceptable type I error rate. Deterministic models of TE load evolution thus have little or no support. Instead the variation of TE loads among the extant species is consistent with a stochastic model. Exclusion of this wide range of direct possible deterministic explanations for TE load variation means that complex interactions among such forces must be postulated to retain strong effect for these proposed mechanisms. This is not to say that such deterministic effects are entirely absent, only that they are short lived due to purifying selection, and disrupted by drift. Drift has been suggested to be key in the evolution of multicellular organisms due to the long-term and ancient reduction in effective population size [91].

### Conclusions

A wide body of literature has sought biological explanations for the observed patterns of variation of TE loads in eukaryotic genomes, invoking explanatory variables such purifying selection, mating system, parasitiic lifestyle, genome-wide GC content and RNAi pathways for silencing TEs. Our analysis of the evolution of TEs on a long time scale – across the entire phylum Nematoda – reveals that purifying selection is strong enough to counteract all other forces, given time. Variation in TE loads on shorter timescales is largely explained by genetic drift, with little or no consistent effect of life history or genomic explanatory variables. We suggest that only studies that examine TE load across a large number of life history transitions and over large timescales will be able to provide power to reliably distinguish between stochastic and deterministic forces, and quantify the balance of evolutionary processes shaping this major component of eukaryotic genomes.

## Materials and Methods

### Genome assemblies

Genome assemblies of species from phylum Nematoda, representing the five major clades [78] were obtained from different sources (S3 Table). The assemblies included four species from Dorylaimida (Clade I), one from Enoplia (Clade II), nine from Spirurina (Clade III), fifteen from Tylenchina (Clade IV) and thirteen from Rhabditina (Clade V). For Dorylaimia and Enoplida (Clades I and II) we analysed all the available genome assemblies. The genome of the tardigrade *Hypsibius dujardini* was used as outgroup. To compare the completeness of the genome assemblies, the N50 metric (S1 Methods, section 1) was calculated for each (S3 Table). The GC content of each genome assembly was calculated (S1 Methods, section 1).

### TE identification

We conducted TE searches in the genome assemblies rather than in sequence read data, which are not publicly available for many of the target species. To mitigate the biases associated with this approach, we have also utilized complementary methods of TE searches. One of the approaches was homology based searches using reference DNA sequences of elements in a *de-novo* constructed library, representing a wide taxonomic range within phylum Nematoda. RepeatModeler 1.0.4 [92] was used to identify repeat sequences in each genome assembly using RECON [93], RepeatScout [94] and TRF [95]. RepeatModeler uses RepeatMasker [96] to classify the consensus sequences of the recovered repetitive sequence clusters. The identification stage employed RMBlast [97] and the Eukaryota TE library from Repbase Update [98]. The consensus sequences from all the species were pooled, and the uclust algorithm in USEARCH [99] was used to make a nonredundant library, picking one representative sequence for each 80% identical cluster. Additional classification of the consensus sequences was performed with the online version of Censor [100]. Classifications supported by matches with a score value larger than 300 and 80% identity were retained. The script used to construct this library is in S1 Methods, sections 2–3.

RepeatMasker [96] was used to search for repeat sequences in the Nematoda genome assemblies and that of *H. dujardini*, using this *de-novo* Nematoda library (S1 Methods, section 4). To eliminate redundancies in RepeatMasker output, we used One Code to Find Them All [101], which assembled overlapping matches with similar classifications, and retained only the highest scoring match of any remaining group of overlapping matches (S1 Methods, section 5). Alternative approaches to identify TEs were also employed. TransposonPSI B (http://transposonpsi.sourceforge.net/), which searches for protein sequence matches in a protein database thus allowing acurate identification of shorter fragments, and LTRharvest [102], which identifies secondary structures (S1 Methods, section 6), were used to screen the target genomes. For TransposonPSI searches, only chains with a combined score larger than 80 were retained, while we retained only matches that were at least 2000 bp long and 80% similar to the query from LTRharvest searches. Where matches from the three approaches overlapped, we retained only the longest match (S1 Methods, section 7).

### Characterization of RNAi pathways

Three key proteins, distinguishing the three RNAi pathways discussed in Sarkies et al. [76], were searched for in the genome assemblies (S1 Methods, section 8.1), using the program Exonerate [103]. Sequences from the supplementary files Data S1 and S2 from Sarkies et al. [76] were used as queries to identify homologues of PIWI, an Argonaute (AGO) subtype, and RNA-dependent RNA polymerase (RdRP; specifically subtypes RRF1 and RRF3) respectively. Only matches at least 100 amino acids (aa) long and at least 60% similar to the query were retained. In addition, only the best scoring out of several overlapping matches was used (S1 Methods, section 8.2). The matches and the queries were used to build two phylogenetic trees, one of PIWI (and other AGO) sequences and the other of RdRP sequences, to verify the identity of the matches (S1 Methods, section 8.3). Each of the datasets was aligned using the L-ins-i algorithm in MAFFT 7 [104], and cleared of positions with a missing data proportion of over 0.3 using trimAl 1 [105]. In the resulting alignment, only sequences longer than 60 aa were retained. Maximum Likelihood (ML) trees were reconstructed using RAxML 8 [106] with sh-like branch supports. Species that occurred at least once in any of the three clades (PIWI in the first tree and RRF1 and RRF3 in the second), were scored as possessing that gene (Figure 1). Where a species did not have a representative sequence in one of the clades, a directed search for the specific protein was conducted in the sequences that did not pass the filter (i.e., the best match had lower score and length than the set cutoff). The identity of sequences retrieved in this way was examined in a second pass of phylogenetic reconstruction. This step did not yield additional phylogenetically validated matches and confirmed the validity of the cutoff set in the filtering step.

### Phylogenetic reconstruction of the Nematoda using small subunit ribosomal RNA (SSU-rRNA) sequences

To control for phylogenetic relationships within the TE counts dataset we inferred a species phylogenetic tree using the SSU-rRNA gene. This locus is considered to be reliable for the reconstruction of the phylogeny of Nematoda, and produces trees that tend to agree with previous analyses [78,107–109]. First, we identified SSU-rRNA genes with BLAST+ 2.2.28 [97], in each of the genome assemblies, where for each species the query was an SSU-rRNA sequence of the same or closely related species, taken from the Silva 122 database [110]. Matches shorter than 1,400 bp were not selected and the query sequence was retained instead, providing it was identical to the match. Species for which the SSU-rRNA sequence could not be recovered and was not available online were excluded from further analysis. Since unbalanced taxon sampling may reduce the accuracy of the phylogenetic reconstruction [111], we also included additional sequences from Silva [110], representing the diversity of Nematoda. ReproPhylo 0.1 [112] was used to ensure the reproducibility of the phylogenetic workflow (S1 Results). A secondary structure aware sequence alignment was conducted using SINA 1.2 [113], and the alignment was then trimmed with trimAl 1 [105] to exclude positions with missing data levels that lie above a heuristically determined cutoff. An ML search was conducted with RAxML 8 [106] under the GTR-GAMMA model and starting with 50 randomized maximum parsimony trees. Branch support values were calculated from 100 thorough bootstrap tree replications. After tree reconstruction, nodes that did not represent a genome assembly (either the blast match or the Silva sequence substitute) were removed from the tree programmatically using ETE2 [114]. To characterize the phylogenetic uncertainty, we generated a posterior distribution of trees using Phylobayes 3 [115]. Two chains were computed, using the trimmed ML tree as a starting tree and the GTR – CAT model *(sensu* [116]). The analysis was continued until the termination criteria were met (specifically, maxdiff and rel_diff < 0.1, and effsize > 100), with a burnin fraction of 0.2 and by sampling each 10th tree. The same subsample of trees was used to generate a consensus tree. The reconstruction of the SSU rRNA tree is detailed in S1 Methods, section 1.

### The effect of life cycle, RNAi and percent GC variation on TE loads

Primary literature was surveyed to determine the mating system of each species and to identify parasites of plants and animals (S1 Table). The effect of these factors on the TE loads was tested with an ANOVA of phylogenetically independent contrasts, using the R package Phytools [117]. Species were classified into the four mating systems dioecy, androdioecy, facultative parthenogenesis (including both species that fuse sister gametes and species that duplicate the genome in the gametes) and strict apomixis. Species that had both hermaphroditic and gonochoric life cycle stages were classified as gonochoric (e.g. *Heterorhabditis bacteriophora*, [118]). We conducted three tests, in the first of which the four levels were tested, in the second the parthenogenetic and androdioecious species were pooled, and in the third, species were divided into dioecious and non-dioecious.

To test the effect of parasitism, free living species, plant parasites and animal parasites were first tested as three separate groups, and then plant and animal parasites were pooled into a single group for a second test. The necromenic lifestyle of *Pristionchus pacificus* was classified as free living because this species is not reported to depend on any host function, only on the organisms that build up on its carcass [119].

ANOVA of phylogenetically independent contrasts was also used to test the effect of the variation in RNAi pathways on the TE loads. Six groups of species were determined based on the presence or absence of PIWI, RRF1 and RRF3 proteins. In addition, for each of the three proteins, the effect of their presence was tested independently of the other proteins. Finally, dependency between GC content of genome assemblies and their TE loads was tested by a regression of the squared contrasts of TE counts and the estimates of GC contents in ancestral nodes [117]. The execution of ANOVA and correlation tests is detailed in S1 Methods sections 10.23–10.25.

### Phylogenetic signal in the TE data

The phylogenetic transformations λ, *κ* and *δ* [79] were calculated with BayesTraits [81] over the subsample of trees produced with Phylobayes (see above), to account for phylogenetic uncertainty (S1 Methods, sections 10.4–10.10). They were estimated for the pooled classes of DNA, LTR, LINE and SINE elements, as well as for individual superfamilies that occurred in at least 15 nematode species.

### Detection of selection and convergent evolution of TE loads

The Ornstein Uhlenbeck (OU) process [120] was originally suggested as an approach to model the evolution of continuous traits based on phylogenies [121]. Building upon this process, Hansen [122] has developed a method to study changes in selection regimes, on the macroevolutionary scale, neglecting stochastic effects on the process. In the OU process, a change in character state depends on the strength of selection (*α*) and its distance and direction from the current selection optimum (*θ*). Goodwin later [123] added a Brownian Motion (BM) component to the model (*σ*^2^), recognizing the confounding effect that stochastic events related to demography might have on selection. The R package PMC [124] was used to assess the power of our data to detect OU processes in the evolution of TEs. The OU parameter *α* was estimated with Bayestraits [81], neglecting stochastic effects, in superfamilies occurring in at least 15 nematode species. Where a significant *α* was detected (p-value < 0.05, in the posterior distribution of trees), indicating selection, we examined the possibility of convergent evolution between species with similar life cycle or RNAi status with the R package SURFACE [125]. In these analyses, selection optima shifts are detected in the trees’ branches through a heuristic search, which uses AIC test results as the optimization criterion. Then, further improvement to the fit of the model is attempted by unifying optimum shifts. Where the unification of two or more optimum shifts improved the AIC score of the model, convergent evolution is inferred. SURFACE uses the R package OUCH [126] to fit OU models, and unlike Bayestraits, includes a stochastic component (*σ*^2^), expressed by BM, in the OU model. The steps described in this paragraph are detailed in S1 Methods section 10.12–10.22.

### Magnitude of change at ancestral nodes

To identify nodes in the species tree that were hotspots of change in TE loads, we reconstructed the ancestral character states for a subset of elements using an ML analysis [117]. The element subset included only classified elements from superfamilies that occurred in at least 15 species. Within each of the three groups of “cut and paste”, LTR, LINE, and SINE elements, we calculated the median change magnitude across the superfamilies in the group, and for each node. The magnitude of change was expressed as the proportion of the load of a given element superfamily at node X out of the load of the same superfamily at the parent of node X. The steps described here are detailed in S1 Methods section 10.23, 10.26–10.27.

## Supplementary Information

**S1 Table:** Life history information

**S2 Table.** – Transposable element counts in Nematoda genome assemblies

**S3 Table:** Genome assemblies used in this study

**S1 Figure:** Transposable element loads in Nematoda genomes

**S2 Figure:** Phylogenetic transformations of Nematoda transposable element counts

**S3 Figure:** Distribution of alpha in a deterministic OU model

**S4 Figure:** Median change in selective optima

**S1 Methods:** Static Jupyter notebooks containing the code used to carry out the analyses in this manuscript

**S1 Results:** Phylogenetic analysis report

## Acknowledgement

We thank Dr. Beth Hellen for her valuable comments, and Dr. Peter Sarkies, Dr. Arvid Ågren and Prof. Carl Boettiger for assistance with the analysis. The following funding sources supported this study: The Science of the Environment Council grant (http://www.nerc.ac.uk/) NE/J011355/1 was awarded to DHL and MLB. The Science of the Environment Council grant (http://www.nerc.ac.uk/) R8/H10/56 was awarded to GenPool, University of Edinburgh. The Medical Research Council grant (http://www.mrc.ac.uk/) G0900740 was awarded to GenPool, University of Edinburgh. The funders had no role in study design, data collection and analysis, decision to publish, or preparation of the manuscript.

